# Effects of autozygosity and schizophrenia polygenic risk on cognitive and brain developmental trajectories

**DOI:** 10.1101/159939

**Authors:** Aldo Cordova-Palomera, Tobias Kaufmann, Francesco Bettella, Yunpeng Wang, Nhat Trung Doan, Dennis van der Meer, Dag Alnæs, Jaroslav Rokicki, Torgeir Moberget, Ida Elken Sønderby, Ole A. Andreassen, Lars T. Westlye

**Affiliations:** NORMENT, KG Jebsen Centre for Psychosis Research, Division of Mental Health and Addiction, Oslo University Hospital & Institute of Clinical Medicine, University of Oslo, Norway; Department of Psychology, University of Oslo, Oslo, Norway

**Keywords:** Neurodevelopment, autozygosity, polygenic risk, cognition, schizophrenia, MRI

## Abstract

Cognitive and brain development are determined by dynamic interactions between genes and environment across the lifespan. Aside from marker-by-marker analyses of polymorphisms, biologically meaningful features of the whole-genome (derived from the combined effect of individual markers) have been postulated to inform on human phenotypes including cognitive traits and their underlying biological substrate.

Here, estimates of inbreeding and genetic susceptibility for schizophrenia calculated from genome-wide data –runs of homozygosity (ROH) and schizophrenia polygenic risk score (PGRS)– are analyzed in relation to cognitive abilities (n=4183) and brain structure (n=516) in a general-population sample of European-ancestry participants aged 8-22, from the Philadelphia Neurodevelopmental Cohort.

The findings suggest that a higher ROH burden and higher schizophrenia PGRS are associated with higher intelligence. Cognition~ROH and cognition~PGRS associations obtained in this cohort may respectively evidence that assortative mating influences intelligence, and that individuals with high schizophrenia genetic risk who do not transition to disease status are cognitively resilient.

Neuroanatomical data showed that the effects of schizophrenia PGRS on cognition could be modulated by brain structure, although larger imaging datasets are needed to accurately disentangle the underlying neural mechanisms linking IQ with both inbreeding and the genetic burden for schizophrenia.

## INTRODUCTION

Cognitive abilities and neural development are determined by complex and dynamic interactions between environmental influences and the individual genetic architecture across the lifespan^1,2^. The genetically-driven temporal regulation is especially noticeable in the transition from childhood to adolescence and adulthood, and can have both immediate and lagged effects on the risk for psychiatric disorders depending on the timing of gene expression and the relevant environmental perturbations, which jointly determine the individual age of onset of the clinical phenotypes^3^. Studies on the genetic architecture of cognitive and structural brain phenotypes are eliciting previous findings from quantitative genetics (i.e., heritability)^4,5^. However, a large fraction of the underlying molecular genetic mechanisms –and their relation to developmental trajectories– remains to be revealed.

Previous research suggests that runs of homozygosity (ROH) are linked to human cognitive abilities^6-8^. Studies on ROH and cognition in different strata of the general population have shown mixed results, with moderate effect sizes^6-8^. While two studies^6,8^ suggest that inbreeding depression may decrease adult intelligence, divergent results by Power et al.^7^ could be explained in view of three evidences. First, assortative mating has recently been highlighted as an important factor underlying psychiatric and behavioral phenotypes^9-12^, in line with specific findings showing assortative mating in relation to cognitive ability published in the late 20^th^ century^13,14^. Secondly, similar to the age-varying pattern for the heritability of cognition^15,16^, the genetic effect of ROH on cognition may be different across the lifespan. Thirdly, differences in ROH burden calculation between the report by Joshi et al.^6^ and the other two studies^7,8^ may have added to the discrepancies.

Growing evidence indicates that the genetic architectures of schizophrenia and cognition are partly shared in young and middle-aged adults^17^, and that the shared genetic variance between cognition and schizophrenia risk might be age-dependent^18^, calling for further research on lifespan patterns. In fact, Germine et al.^19^ reported that higher schizophrenia PGRS is linked to lower speed of emotion identification and speed of verbal reasoning in healthy young individuals from the Philadelphia Neurodevelopmental Cohort (PNC). Consistently, Shafee et al.^20^ recently found an association between schizophrenia PGRS and specific cognitive abilities in adults. The latter findings may be mediated by genetic effects on brain structure and function, but the evidence of associations between genetic risk for schizophrenia and brain features is scarce, with some studies reporting no overlap between schizophrenia PGRS and subcortical phenotypes derived from magnetic resonance imaging (MRI)^21^, whereas others suggest an association with cortical thickness/surface area^22^. Replication studies using independent samples and probably a wider span of brain features is thus needed.

With this background, using a sample of 4183 young participants (ages 8-22) with European-ancestry from the PNC, we pursue the following four main aims. First, the putative links between multivariate genome-wide features (ROH burden and schizophrenia PGRS, computed independently) and both intellectual abilities and neuroanatomical features are tested. Secondly, potential age-modulating effects on relevant outcomes of the latter analysis are tested. Thirdly, whether inbreeding modifies the effects of genome-wide schizophrenia burden on cognition and brain anatomy is evaluated by testing the effects of the interaction between ROH and PGRS on cognitive ability and brain structure. Finally, the presence of brain anatomy differences mediating the associations between ROH/PGRS and cognition is assessed from a causal mediation framework. Using those elements, our overall goal is to outline potential neurodevelopmental pathways leading from biologically meaningful whole-genome features to neural and ultimately cognitive disruptions.

## MATERIALS AND METHODS

### Participants and measures

The data was retrieved from the PNC public domain resources. The PNC includes more than 9,000 individuals aged 8-22 years drawn from a larger population, enrolled through a joint collaborative effort of the Center for Applied Genomics at Children’s Hospital of Philadelphia and the Brain Behavior Laboratory (University of Pennsylvania). All enrolled participants were able to provide informed consent, and parental consent was also required for subjects under age 18. All individuals underwent a psychiatric evaluation using a structured clinical interview, and the Wide Range Achievement Test (WRAT)^23^ was applied. After removing participants with non-European ancestry or due to missing data for relevant variables, 4,183 individuals were included in the main analyses (ROH, PGRS and cognition), and MRI data was available for 516 of them. A more comprehensive description of the PNC demographics and data collection protocols is provided in previous publications^24-26^. PNC data is publicly available via dbGaP (https://www.ncbi.nlm.nih.gov/projects/gap/cgi-bin/study.cgi?study_id=phs000607.v1.p1) and related resources (https://www.med.upenn.edu/bbl/philadelphianeurodevelopmentalcohort.html). Analysis pipelines used in this study can be obtained upon request to the authors.

### Genotyping platforms and genotype imputation

Genotyping was performed by the Center for Applied Genomics, at the Children’s Hospital of Philadelphia. The DNA samples from the PNC were genotyped in different batches using the following platforms: Illumina OmniExpress (*n* = 1,657), Illumina Human-610 Quad (*n* = 3,807), Illumina HumanHap-550-v1 (*n* = 556), Illumina HumanHap-550-v3 (*n* = 1,914), Affymetrix Genome-Wide Human SNP Array 6.0 (*n* = 66) or Affymetrix Axiom (*n* = 722) (hereafter, Omni, Quad, 550-v1, 550-v3, Affy60 and Axiom). From those datasets, only participants with European ancestry were included in the final statistical analyses, in recognition that the inclusion of subjects from other ethnicities might add genetic variability altering both ROH- and PGRS-phenotype association estimates (e.g., homozygosity might differ between ethnicities and across samples with distinct admixture levels^8^, and schizophrenia PGRS might not explain much variance in non-European samples^27^). Further details on genotype imputation can be found in Supplementary Material.

### Runs of homozygosity (ROH)

The sum of the total length of ROHs across chromosomes 1 to 22 was divided by the total SNP-mappable autosomal length (2.77 × 10^−9^ bp), to obtain an estimate of ROH burden for each individual (ROH fraction, *F*_ROH_). The protocol was based on parameters widely used in previously published manuscripts^7,8,28^, with slight modifications based on quality control procedures suggested elsewhere^29^. Briefly, each of the filtered genotyping batches was pruned for LD using VIF threshold = 10, with window size = 50 SNPs and window shift step = 5 SNPs. Then, *F*_ROH_ was calculated with PLINK’s sliding-window approach, with a minimum SNP length threshold and window size of 65 SNPs, 5% of missing SNPs allowed and no heterozygote SNPs accepted. The number of independent ROH segments was also extracted from the “.hom.indiv” files generated by PLINK, and is hereafter referred to as *N*_ROH_. Additional details on the ROH-calling algorithm are reported in Supplementary Material.

### Schizophrenia PGRS

The imputed genotypes passing quality control (see *Genotyping platforms and genotype imputation*) were used to compute schizophrenia PGRS, based on data published by the Schizophrenia Working Group of the Psychiatric Genomics Consortium^30^. Initially, SNPs with ambiguous alleles (AT or CG), or in linkage disequilibrium with the local SNP with the smallest *p*-value were removed (the SNP with the smallest *p*-value within a 250kb window is retained, and all neighbors with a linkage disequilibrium *r*^2^ > 0.1 are removed; a step known as clumping). Also, SNPs within the major histocompatibility complex region (chromosome 6, 26-33Mb) were omitted, and five hundred different PGRS values were obtained (*p*-value thresholds, *p*_threshold_, from 0.001 to 0.5, with intervals of 0.001). The former procedures were implemented in PRSice.

As done in former studies^31-33^, a specific *p*-value cutoff was chosen for the schizophrenia PGRS (out of the five hundred estimations) by tuning the fitting parameters (adjusted *R*^2^) to maximize the explained variance of independent regression models with cognitive performance as outcome and gender, age, batch and PGRS as dependent variables.

### Image acquisition and pre-processing

MRI was performed for a subset of participants at the Hospital of the University of Pennsylvania, by means of a 3T Siemens TIM Trio whole-body scanner (32-channel head coil; gradient performance: 45 mT/m; maximum slew rate: 200 T/m/s). The focus of the current study is on structural brain features derived from 3D T1-weighted images obtained using a MPRAGE sequence (TR: 1.81s; TE: 3.5ms; FA: 9°; FOV: 240×180mm; 1mm slice thickness; 160 slices), whose acquisition parameters have formerly been described in more detail^26^.

Data from each participant were pre-processed using the recon-all stream from Freesurfer v5.3.0 (http://surfer.nmr.mgh.harvard.edu/)^34^, using automatic parcellation and segmentation protocols to obtain 68 cortical and 14 subcortical gray matter brain regions, as well as some global brain features (e.g., intracranial volume (ICV))^35^. Thickness and surface measurements from cortical regions were used for the ensuing analyses, along with volumetric estimates of subcortical volumes. Twelve relevant brain features were selected for analysis: intracranial and seven subcortical volumes used in a recent ENIGMA report^4^, along with mean cortical thickness, total cortical surface area, cerebellar cortex volume and cerebellar white matter volume. After merging with genetic and other phenotypic data of European-ancestry participants, there were 516 participants with MRI measures available. A measure of participants’ age in months was extracted from the DICOM headers of the 516 MRI files, and it was used in the analyses involving brain features.

### Statistical analyses

The analyses of ROH, PGRS and cognition were performed via linear regression in R, including sex, age, age^2^ and ten genetic principal components as covariates. For the analysis of ROH, fraction of missing values and SNP-by-SNP homozygosity were also adjusted for. Finally, the question of whether differences in brain features mediate the association between genomic features and IQ was assessed using causal mediation. Results reported below include *p*-values adjusted using either Bonferroni correction or false discovery rate^36,37^, which have been documented as appropriate methods to control for multiple testing. In the regression models, all continuous variables were scaled with R’s “scale” function, to get interpretable coefficients. Additional details on the statistical methods are included in *Supplementary Material*.

## RESULTS

### Demographic and clinical data

The data shown in Table 1 and Figure 1 summarizes phenotypic information of PNC’s sample subset included herein. When considering all participants, the correlations between age, age^2^ and total WRAT score were either small or moderate, with the largest coefficient for WRAT and age (*r* = 0.67). Even though the PNC participants were initially recruited randomly from the set of available genotypes regardless of genotyping platform^26^, the age and IQ distributions displayed small yet statistically significant differences across batches (median ages: 13, 14, 15 and 16 years, *p*<3×10^−16^; median IQ scores: 105 and 106, *p*=0.006), which may have had an impact on the distribution of other variables such as ICV (*p*=0.024 for between-batch differences) and related neuroimaging measures. Besides, the contingency table of medical status categories (none/minor, mild, moderate and severe) and genotyping platforms displayed unbalanced frequencies (*p*=10^−10^, Table 1). These observations, along with the differences in genetic measures discussed below, encouraged the inclusion of genotyping platform in all ensuing analyses.

**Figure 1.**
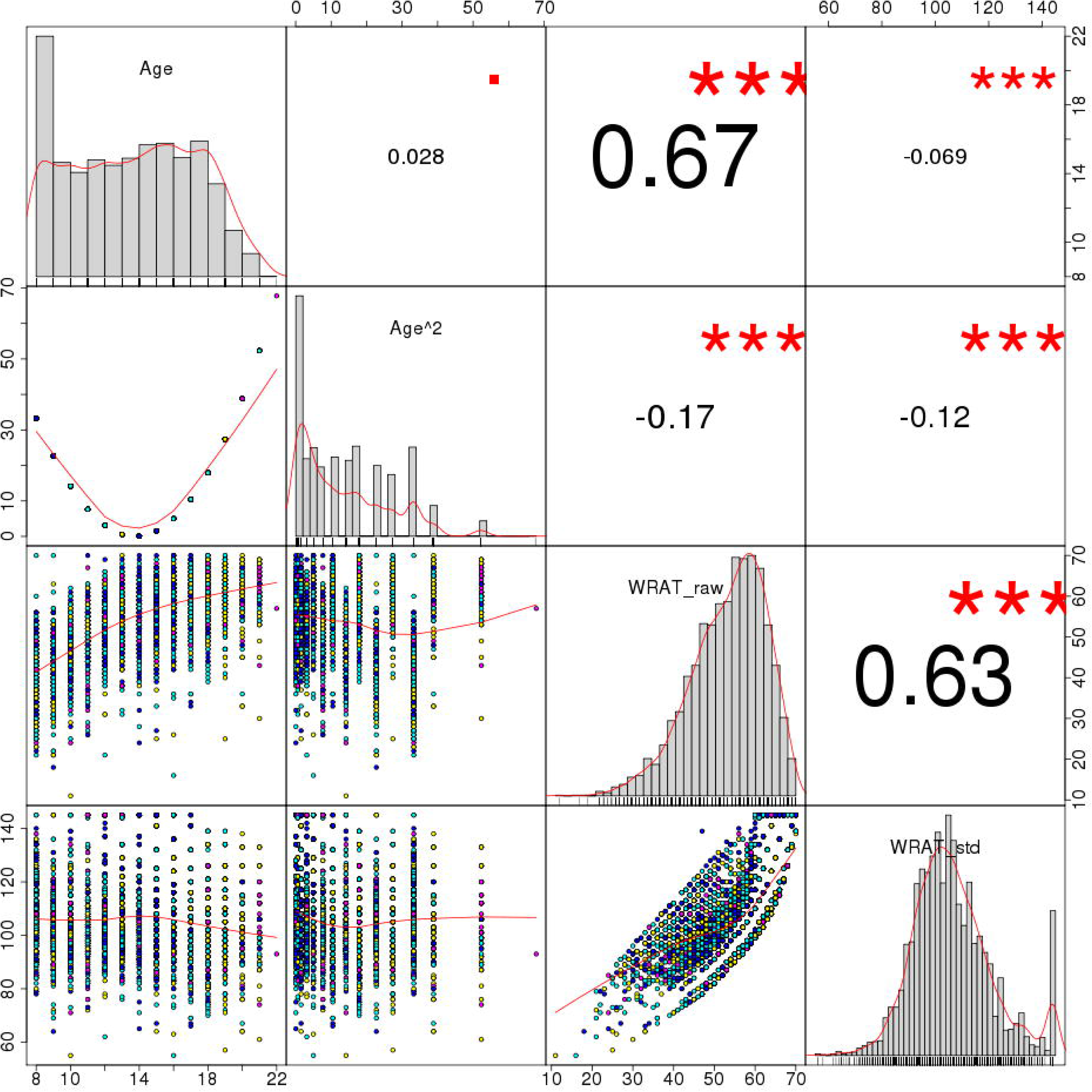
Distribution of age, age squared, total WRAT scores and age-standardized WRAT scores Notes: age range in the uppermost left square is shown in years (8-22), and WRAT scores are displayed both before (raw) and after (std) age standardization. Numbers on the upper right triangle correspond to Pearson correlation coefficients between pairs of variables, and symbols indicate significance levels (■: *p*<0.1, *: *p*<0.05, **: *p*<0.01, ***: *p*<0.001).

**Table 1.** Descriptive values of the main phenotypic and genetic variables included in the analyses, stratified by genotyping platform Notes: a, age at Computerized Neurocognitive Battery test date; b, values of estimated total intracranial volume were available only for a subset of participants (*n* = 385) with genetic and cognitive information included in the study (see *Materials and methods – Image acquisition and pre-processing*); c, schizophrenia PGRS at the best-fitting *p*_threshold_ (0.005), as mentioned in the manuscript; d, between-group comparisons were performed using analysis of variance (ANOVA) tables for most of the variables displayed (continuous measures; *F* and *p* values reported), except for gender and medical status, where chi-squared tests were applied on the contingency tables.

|  |  | Omni | Quad | 550-v1 | 550-v3 | Between-group comparison statistic (*p*-val.)<sup>d</sup> |
| --- | --- | --- | --- | --- | --- | --- |
| Subjects (*n*) | male/female | 573/590 | 886/896 | 148/154 | 462/474 | 0.094 (0.993) |
| Medical status | 0 / 1 / 2 / 3 / 4 | 207 / 321 / 342 / 213 / 78 | 264 / 529 / 496 / 368 / 112 | 30 / 56 / 105 / 81 / 30 | 103 / 220 / 337 / 216 / 49 | 72.804 (10<sup>-10</sup>) |
| Age at CNB<sup>a</sup> (years) | Median | 13 | 14 | 16 | 15 | 30.922 (<3×10<sup>-16</sup>) |
|  | Range | 8 – 21 | 8 – 21 | 8 – 22 | 8 – 21 |  |
| Raw WRAT score | Median | 54 | 55 | 57 | 55 | 7.091 (9.5×10<sup>-5</sup>) |
|  | Range | 18 – 70 | 16 – 70 | 24 – 69 | 11 – 70 |  |
| Standardized WRAT score (IQ, standard units) | Median | 106 | 105 | 106 | 105 | 4.106 (0.006) |
|  | Range | 64 – 145 | 55 – 145 | 71 – 145 | 55 – 145 |  |
| Intracranial volume (10<sup>6</sup> mm<sup>3</sup>)<sup>b</sup> | Median | 1.609 | 1.603 | 1.55 | 1.573 | 2.292 (0.077) |
|  | Range | 1.218 – 1.988 | 1.095 – 1.98 | 1.253 – 1.893 | 1.179 – 1.849 |  |
| PGRS<sup>c</sup> | Median | -3.935×10<sup>-3</sup> | -3.926×10<sup>-3</sup> | -3.962×10<sup>-3</sup> | -3.952×10<sup>-3</sup> | 7.238 (8×10<sup>-5</sup>) |
|  | Range | -4.397×10<sup>-3</sup> – -3.509×10<sup>-3</sup> | -4.45×10<sup>-3</sup> – -3.386×10<sup>-3</sup> | -4.363×10<sup>-3</sup> – -3.463×10<sup>-3</sup> | -4.427×10<sup>-3</sup> – -3.422×10<sup>-3</sup> |  |
| *F*<sub>SNP</sub> | Median | 0.6×10<sup>-3</sup> | 0.001 | -0.9×10<sup>-3</sup> | 0.7×10<sup>-3</sup> | 15.315 (7×10<sup>-10</sup>) |
|  | Range | -0.017 – 0.039 | -0.023 – 0.133 | -0.019 – 0.012 | -0.018 – 0.052 |  |
| *F*<sub>miss</sub> | Median | 0.5×10<sup>-3</sup> | 93×10<sup>-6</sup> | 0.1×10<sup>-3</sup> | 0.2×10<sup>-3</sup> | 74.885 (<3×10<sup>-16</sup>) |
|  | Range | 0.1×10<sup>-3</sup> – 5×10<sup>-3</sup> | 12×10<sup>-6</sup> – 0.01 | 0.02×10<sup>-3</sup> – 8.9×10<sup>-3</sup> | 0.01×10<sup>-3</sup> – 12.8×10<sup>-3</sup> |  |
| *N*<sub>ROH</sub> | Median | 3 | 3 | 2 | 3 | 7.63 (4×10<sup>-5</sup>) |
|  | Range | 0 – 19 | 0 – 40 | 0 – 12 | 0 – 21 |  |
| *F*<sub>ROH</sub> | Median | 1.652×10<sup>-3</sup> | 1.603×10<sup>-3</sup> | 1.331×10<sup>-3</sup> | 1.438×10<sup>-3</sup> | 1.659 (0.174) |
|  | Range | 0 – 41×10<sup>-3</sup> | 0 – 127×10<sup>-3</sup> | 0 – 12×10<sup>-3</sup> | 0 – 49×10<sup>-3</sup> |  |

### Schizophrenia PGRS and cognitive performance

As mentioned above (see *Methods*) five hundred schizophrenia PGRS values were computed (to assess the *p*-value threshold (*p*_threshold_) range from 0.001 to 0.5, in steps of 0.001). The model fitting parameters of PGRS for IQ are shown in Supplementary Figure S1, using four different settings: models with or without participants displaying moderate/severe medical conditions, evaluated using either 4 or 15 covariates along with the PGRS. Outcomes were assessed based on both the overall adjusted-*R*^2^ of the full model, and the PGRS’s *p*-value within that model. Data on Supplementary Figure S1 provide multiple insights: first, *15-covariate* models showed higher adjusted-*R*^2^ values than their *4-covariate* counterparts, suggesting that the principal components extracted from the identity-by-state do explain part of the phenotypic variance in cognitive performance. Second, schizophrenia PGRS display smaller *p*-values when including participants with medical conditions. Third, across all models, the best fitting parameters (smallest *p*-values) throughout the different threshold values were observed using a *p*_threshold_ of either 0.005 or 0.01. Of note, when using standardized WRAT scores (a measure of IQ), 0.005 was also the threshold with the smallest *p*-values; it was thus selected for the analyses shown next (Supplementary Figure S1 and Supplementary Figure S2). The results indicate that a higher schizophrenia PGRS is associated with higher cognitive performance across age (β=0.027, SE=0.011, *p*=0.015; Supplementary Figure S3); β values were positive across PGRS *p*-value thresholds in this model (range: 0.02-0.35), as well as in all *4-covariate* regressions and excluding participants with moderate/severe medical conditions. Lastly, there were no significant interactions between PGRS and either age or age^2^ on cognitive performance in the full dataset (*n*=4,183; PGRS×age: β=-0.02, SE=0.011, *p*=0.071; PGRS×age^2^: β=0.014, SE=0.011, *p*=0.21), although there was an association with age when excluding participants with medical conditions (*n*=3,036; PGRS×age: β=-0.026, SE=0.013, *p*=0.037; PGRS×age^2^: β=0.019, SE=0.013, *p*=0.139; Supplementary Figure S4). Those results were similar when using *4-covariate* models.

### ROH and cognitive performance

Descriptive values of *F*_ROH_ and related variables (*N*_ROH_, *F*_SNP_, *F*_miss_) are shown in Table 1 and Figure 2. There were no significant differences in the distribution of *F*_ROH_ across platforms, although a statistically significant batch-specific pattern was observed in *F*_miss_, *F*_SNP_ and *N*_ROH_ (*p*<3×10^−16^, *p*=7×10^−10^ and *p*=4×10^−5^). Pairwise analysis of these variables indicated a moderate correlation between the number of ROH per individual (*N*_ROH_) and SNP-by-SNP homozygosity (*F*_SNP_) (*r*=0.38), whereas larger coefficients were observed between *F*_ROH_ and *F*_SNP_ (*r*=0.5), and between *F*_ROH_ and *N*_ROH_ (*r*=0.75). The latter is shown in more detail in Supplementary Figure S5, where the *F*_ROH_ and *N*_ROH_ values are compared for the whole sample. The correlations remained virtually unchanged after removing participants with moderate/severe medical status. Those two distributions (with and without participants with relevant medical conditions) are highly overlapping, with only two out of six high-ROH burden participants (*F*_ROH_>0.025) coming from the subsets of moderate and severe medical conditions. Of note, *F*_ROH_ was linearly independent of the schizophrenia PGRS (full sample: *r*=0.016, *p*=0.293; removing participants with medical conditions: *r*=0.025, *p*=0.172) (Figure 2 and Supplementary Figure S5), which further justifies the analysis of their joint effects in subsequent tests.

Regarding phenotype-ROH burden association tests, the output of linear regression analysis in the whole dataset (*n*=4,183) showed a positive-signed association between cognitive performance and *F*_ROH_ (β=0.047, SE=0.013, *p*=3.5×10^−4^, Supplementary Figure S6). Despite the apparent bias introduced by a few subjects with very high *F*_ROH_ values within the *Quad* platform, the effect was noticed across the four genotyping batches (Supplementary Figure S7), and the analysis excluding subjects with *F*_ROH_>0.02 displayed similar outcomes (β=0.057, SE=0.012, *p*=1.3×10^−6^). The results after excluding participants with moderate/severe medical conditions (namely, keeping 3,036 individuals) remained statistically significant (β=0.046, SE=0.014, *p*=1.6×10^−3^). An additional analysis revealed no interactions between *F*_ROH_ and either age or age^2^ on IQ (*F*_ROH_×age: β=0.011, SE=0.011, *p*=0.325; *F*_ROH_×age^2^: β=-0.007, SE=0.016, *p*=0.672; Supplementary Figure S8).

**Figure 2.**
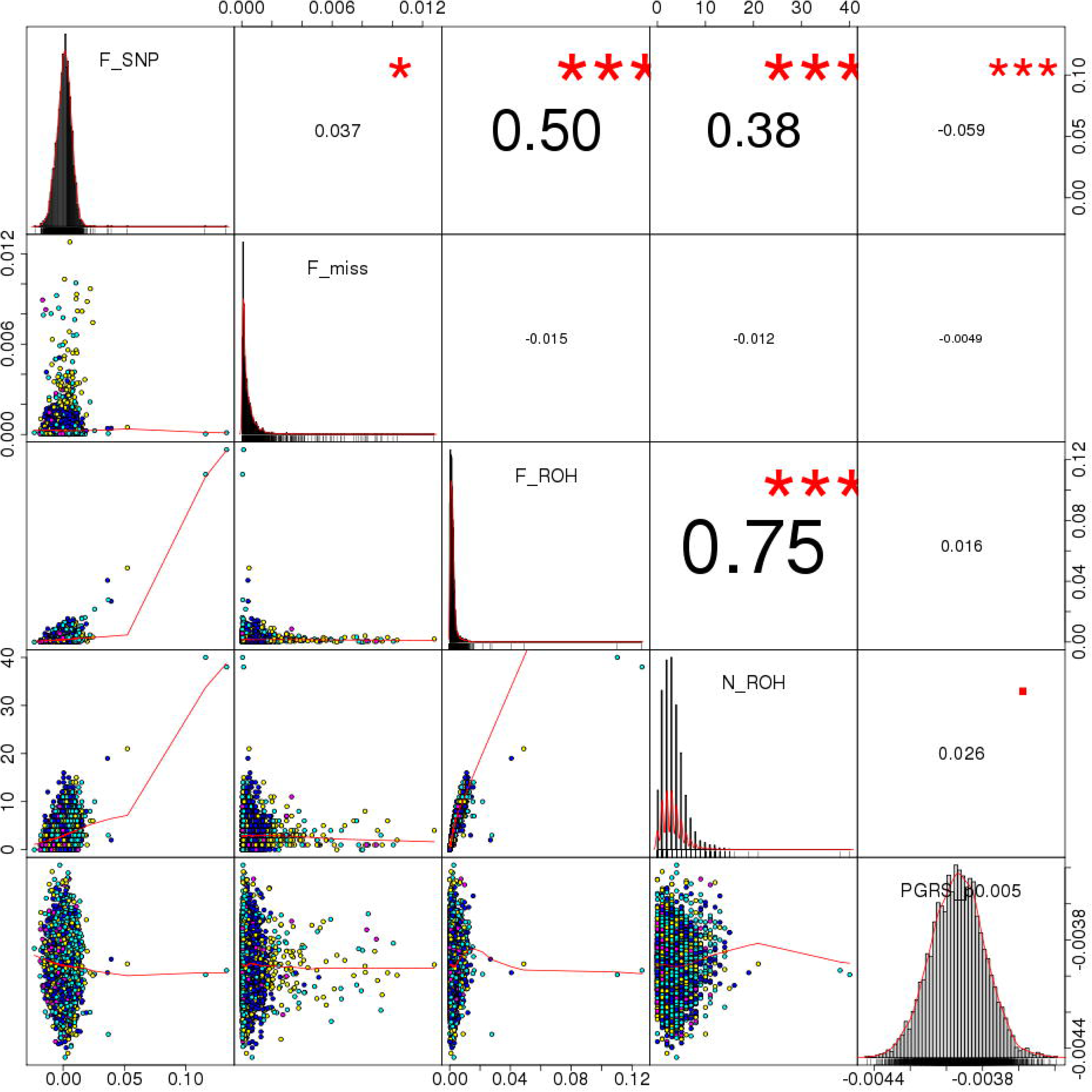
Correlation between different genomic features Notes: F_SNP represents the SNP-by-SNP homozygosity; F_miss indicates the fraction of missing genotypes per sample; F_ROH is the fraction of the genome covered by homozygous blocks (included as variable of interest in the downstream analysis); N_ROH corresponds to the number of homozygous segments of each participant; PGRS_p0.005 denotes schizophrenia polygenic risk scores using markers with *p*≤0.005 in the PGC2 GWAS. Numbers on the upper right triangle correspond to Pearson correlation coefficients between pairs of variables, and symbols indicate significance levels (■: *p*<0.1, *: *p*<0.05, **: *p*<0.01, ***: *p*<0.001).

### ROH × schizophrenia PGRS interaction on IQ

A statistically significant interaction effect between ROH burden and schizophrenia PGRS on cognitive performance was detected, although the explained variance was relatively small. Briefly, the main effects model (*15-covariate*) showed significant results for both *F*_ROH_ (β=0.046, SE=0.013, *p*=5.2×10^−4^) and schizophrenia PGRS (β=0.025, SE=0.011, *p*=0.023), with an adjusted-*R*^2^ of 0.4864 for the whole regression, whereas in the multiplicative interaction model *F*_ROH_ × PGRS was statistically significant (β=0.032, SE=0.016, *p*=0.048; overall adjusted-*R*^2^: 0.4867). As indicated, there was slight increase in the model fitting parameter (adjusted *R*^2^ shifting from 0.4864 to 0.4867), with a statistically significant effect according to the ANOVA test for the interaction (*F*=3.9, *p*=0.048). The results were very similar within the *4-covariate* framework, with significant effects for both *F*_ROH_ and PGRS (*F*_ROH_: β=0.037, SE=0.011, *p*=7.8×10^−4^; PGRS: β=0.027, SE=0.011, *p*=0.015; overall adjusted-*R*^2^: 0.4861), with a significant interaction term and improved model-fitting statistics (*F*_ROH_×PGRS: β=0.036, SE=0.016, *p*=0.029; overall adjusted-*R*^2^: 0.4866), and statistically significant ANOVA results (*F*=4.8, *p*=0.029). The data in Supplementary Figure S9 indicates that individuals with a high level of inbreeding would be more sensitive to the effects of schizophrenia polygenic burden: those in the uppermost *F*_ROH_ quartile would have higher cognitive performance when their schizophrenia PGRS is high. In contrast, subjects with lower *F*_ROH_ generally display no correlation between PGRS and cognition. As evidenced in Supplementary Figure S9, despite statistically significant, the effect sizes of these interactions were relatively small. Of note, when excluding participants with high homozygosity burden (*F*_ROH_>0.02), the main effects showed similar effect sizes and significance levels (*F*_ROH_: β=0.055, SE=0.012, *p*=2.5×10^−6^; schizophrenia PGRS: β=0.024, SE=0.011, *p*=0.032), but the interaction term was not statistically significant (β=0.009, SE=0.011, *p*=0.422). There were no three-way interactions with age or age^2^ (*F*_ROH_×PGRS×age: β=-0.019, SE=0.016, *p*=0.235; *F*_ROH_×PGRS×age^2^: β=-0.001, SE=0.019, *p*=0.951). The significance of the findings remained unchanged after removing participants based on medical status.

### Brain features, ROH and schizophrenia PGRS

Descriptive values of the brain features, in relation to age and cognitive performance (age-standardized) are shown in Figure 3 and Supplementary Figure S10, respectively. Table 2 displays the results of *15-covariate* models evaluating the associations either between brain features and *F*_ROH_ or between brain features and schizophrenia PGRS. After multiple testing adjustments, higher PGRS was significantly associated with smaller cortical surface area and smaller thalamic volume. Although there was no association between PGRS and cognitive ability in the subset of participants with MRI data (*n*=516), causal mediation analysis suggested a suppression effect: modifications of brain feature sizes would compensate against the direct effect of schizophrenia PGRS on cognitive ability (cortical area: Average Causal Mediation Effects (ACME)=-0.026, 95% C.I.=[−0.059, - 0.002], *p*=0.028, Average Direct Effect (ADE)=0.058, 95% C.I.=[−0.115, 0.232], *p*=0.52; thalamus: ACME=-0.021, 95% C.I.=[−0.051, −0.0004], *p*=0.045, ADE=0.053, 95% C.I.=[−0.119, 0.225], *p*=0.55). Notice that such assertion had only limited support from the mentioned data subset, since no direct effects (ADE) of PGRS were detected.

With regards *F*_ROH_, there was no statistically significant association. PGRS×*F*_ROH_ interaction tests did not reveal any significant association on the assessed brain features.

**Figure 3.**
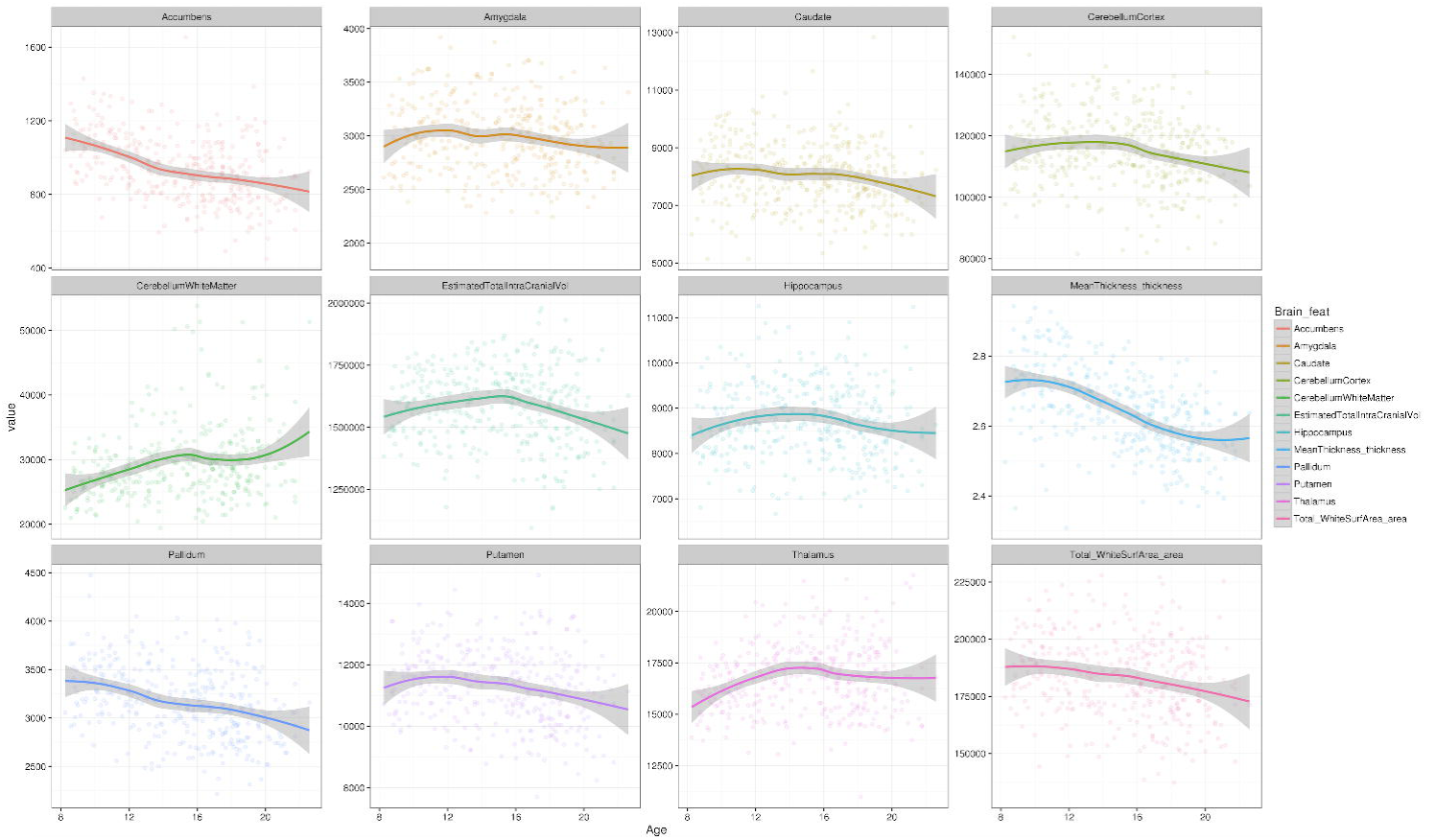
Brain feature values, plotted against age Notes: the vertical scales are in mm^3^ for most features (volumetric traits), except for mean cortical thickness (in mm) and total cortical surface area (in mm^2^). Age (horizontal axis) is displayed in years.

**Table 2.** Association between brain features and either schizophrenia PGRS or *F*_ROH_ Notes: The analyses were performed on 516 participants with both genetic and cognitive information included in the study (see *Materials and methods – Image acquisition and pre-processing*). Results correspond to the *15-covariate* model, and ICV was included as a covariate in tests of subcortical, cerebellar and cortical surface area. FDR: false discovery rate. *, significant *p*-value (less than 0.05) after FDR adjustment.

| | Schizophrenia PGRS | | | | | | $F_{ROH}$ | | | | | |
| --- | --- | --- | --- | --- | --- | --- | --- | --- | --- | --- | --- | --- |
|  | Estimate | SE | t value | p (>\|t\|) | FDR p | Bon-ferroni p | Estimate | SE | t value | p (>\|t\|) | FDR p | Bon-ferroni p |
| **Total_WhiteSurfArea_area** | -8.045E-02 | 2.754E-02 | -2.922E+00 | 3.642E-03 | 2.471E-02 | 4.370E-02 | 1.098E-02 | 5.194E-02 | 2.114E-01 | 8.327E-01 | 9.807E-01 | 1.000E+00 |
| **Accumbens** | 1.189E-01 | 4.126E-02 | 2.882E+00 | 4.118E-03 | 2.471E-02 | 4.941E-02 | 1.347E-01 | 7.757E-02 | 1.736E+00 | 8.319E-02 | 5.386E-01 | 9.982E-01 |
| **Amygdala** | -7.208E-02 | 4.143E-02 | -1.740E+00 | 8.249E-02 | 2.475E-01 | 9.898E-01 | 7.925E-02 | 7.763E-02 | 1.021E+00 | 3.078E-01 | 7.386E-01 | 1.000E+00 |
| **Caudate** | -2.061E-03 | 3.998E-02 | -5.155E-02 | 9.589E-01 | 9.589E-01 | 1.000E+00 | -1.078E-01 | 7.460E-02 | -1.445E+00 | 1.491E-01 | 5.386E-01 | 1.000E+00 |
| **Hippocampus** | -6.505E-02 | 4.091E-02 | -1.590E+00 | 1.125E-01 | 2.699E-01 | 1.000E+00 | -1.111E-01 | 7.653E-02 | -1.452E+00 | 1.471E-01 | 5.386E-01 | 1.000E+00 |
| **Pallidum** | 4.656E-02 | 3.757E-02 | 1.239E+00 | 2.158E-01 | 3.700E-01 | 1.000E+00 | 8.508E-03 | 7.036E-02 | 1.209E-01 | 9.038E-01 | 9.807E-01 | 1.000E+00 |
| **Putamen** | 3.530E-02 | 3.878E-02 | 9.102E-01 | 3.631E-01 | 4.372E-01 | 1.000E+00 | 4.514E-02 | 7.256E-02 | 6.221E-01 | 5.341E-01 | 9.157E-01 | 1.000E+00 |
| **Thalamus** | -6.855E-02 | 3.189E-02 | -2.150E+00 | 3.207E-02 | 1.283E-01 | 3.849E-01 | -8.039E-02 | 5.981E-02 | -1.344E+00 | 1.795E-01 | 5.386E-01 | 1.000E+00 |
| **CerebellumWhiteMatter** | 5.017E-02 | 4.429E-02 | 1.133E+00 | 2.579E-01 | 3.869E-01 | 1.000E+00 | -2.007E-03 | 8.294E-02 | -2.419E-02 | 9.807E-01 | 9.807E-01 | 1.000E+00 |
| **CerebellumCortex** | -3.134E-02 | 4.140E-02 | -7.570E-01 | 4.494E-01 | 4.903E-01 | 1.000E+00 | -6.707E-02 | 7.741E-02 | -8.664E-01 | 3.867E-01 | 7.734E-01 | 1.000E+00 |
| **MeanThickness_thickness** | 3.382E-02 | 3.725E-02 | 9.079E-01 | 3.644E-01 | 4.372E-01 | 1.000E+00 | -2.335E-02 | 6.985E-02 | -3.342E-01 | 7.383E-01 | 9.807E-01 | 1.000E+00 |
| **EstimatedTotalIntraCranialVol** | -5.913E-02 | 4.004E-02 | -1.477E+00 | 1.404E-01 | 2.808E-01 | 1.000E+00 | -1.736E-02 | 7.520E-02 | -2.309E-01 | 8.175E-01 | 9.807E-01 | 1.000E+00 |

## DISCUSSION

In this work, the potential influence of both autozygosity and cumulative genetic risk for schizophrenia on cognitive performance was evaluated in a large and harmonized cohort (*n*=4,183) of European-ancestry participants from the general population, aged 8-22. Increased inbreeding, as indexed by a larger fraction of the genome in ROH, was associated with higher cognitive performance. Similarly, a higher genetic burden for schizophrenia was related to higher cognitive scores. Although ostensibly paradoxical, the results agree with some previous reports using related study designs^7,38,39^. Additionally, interaction effects suggest that more inbred individuals might display higher cognitive test scores in the presence of high schizophrenia polygenic risk score. The relatively small effect sizes indicate that the contribution of these whole-genome features to the total heritability of cognitive performance is modest.

Regarding ROH and cognition, perhaps the most similar study to this one was conducted by Power et al.^7^ in a demographically analogue study sample (2,329 European-ancestry participants, age 12). The ROH-calling procedures using both here and in the former report were also equivalent, and in both cases the outcomes suggest that, in young individuals, increased autozygosity is associated with higher intelligence. To interpret their findings, Power et al.^7^ hypothesized that positive assortative mating in couples with better cognitive profiles could partly explain the mentioned association. Observations from other independent reports support the hypothesis of differential assortative mating patterns influencing cognitive, behavioral and psychiatric phenotypes^9-14^, which is possibly the most plausible framework to explain the findings from Power et al.^7^ and the current report. Importantly, further research is needed since both research outcomes indicate relatively small effect sizes which might have been shadowed by unpublished findings.

Importantly, two former studies have found the inverse association in samples of adult participants. First, Joshi et al.^6^ found a negative correlation between ROH and cognition by meta-analyzing information of 53,300 participants from different cohorts and ancestries. Importantly, as the same authors showed, both *F*_ROH_ and *N*_ROH_ vary depending on ancestry, which might have induced heterogeneity in that study that is not present in the current dataset. It is worth noting that Joshi et al.^6^ included datasets (e.g., FTC_1) that are relatively similar to the present study sample (PNC) in terms of demographics, but there were no statistically significant results within those sub-cohorts. Differences in ROH-calling procedures could have also influenced this between-study discordance. Here, a validated protocol^28^ of increasing popularity has been employed, which could strengthen the reliability of the findings. Another study by Howrigan et al.^8^ showed, in a sample of 4,854 European-ancestry adults from nine cohorts, that increased ROH burden might be associated with lower intelligence. The main focus on adult participants in both previous studies may limit the comparability of the results, since the genetics of intelligence exhibit largely dynamic patterns over the lifetime^15,16^.

Interestingly, we found a positive correlation between schizophrenia PGRS and cognitive performance, suggesting that higher genetic burden for schizophrenia is related to better cognitive performance. This finding somehow agree with observations from two previous cohorts of healthy participants, which indicate that higher schizophrenia PGRS would be associated with decreased risk for psychosis-like experiences and schizotypy^38,39^. The findings from those two reports were interpreted by^38^ in view of potential involuntary biases in sampling: samples with only *healthy* participants are not likely to include subjects with high schizophrenia PGRS and high psychosis-related phenotypes; people with high schizotypy and low resilience would seldom be part of the healthy general population, but would transition to clinical psychosis instead. Within that framework, lower genetic disease risk would protect healthy high-schizotypy individuals against transition to the clinical phenotype. A high-PGRS-low-resilience population subset would then rarely be part of a *healthy* sample, not because of an explicit sampling bias to exclude subclinical phenotypes, but rather due to psychopathological dynamics leading high-PGRS-low-resilience to the *affected/patient* groups. Of note, the current data (PNC) includes participants across a broad spectrum of phenotypes ranging from health to different diseases, and the association between higher schizophrenia PGRS and better cognition is more noticeable when removing participants with medical conditions. Namely, the positive correlation between PGRS and cognitive ability is stronger in the *healthiest* end of the phenotypic distribution, probably in concordance with an interplay between resilience and genetic schizophrenia risk as referred above.

Regarding the latter finding on schizophrenia PGRS and IQ, it is also important to notice that IQ has recently been reported to have a small negative genetic correlation with schizophrenia risk^40^. Those results obtained with LD Score Regression^41^ are ostensibly in disagreement with the current findings, but it is also important to highlight that the data from the Brainstorm Consortium indicate a positive correlation between multiple psychiatric diseases (bipolar disorder, autism and anorexia) and both education years and college attainment, and the absence of genetic correlation between cognitive performance and schizophrenia^42^. In addition, Okbay et al.^43^ have also found a positive genetic correlation between years of education and schizophrenia; overall, there are mixed results that warrant further research to validate whether an increased genetic risk for schizophrenia is linked to the genetics of intelligence. While the arguments on unintended sampling bias outlined before –as opposed to true PGRS causality– are probably the most complete interpretation for the present findings, the latter evidences from other studies could also suggest that schizophrenia PGRS directly increases cognitive scores.

In addition to the explanation above, the observed association between increased genetic risk for schizophrenia and higher cognitive ability might partly be due to the link between psychosis PGRS and psychological traits such as creativity^44^, which might be closely related to intelligence^45^. As discussed in the literature on psychometrics, there seem to be discontinuous populations when comparing IQ and creativity: creativity would be higher in individuals above a certain IQ score^45^; analogous non-steady relationships might also be postulated when stratifying IQ by genetic risk for psychosis: perhaps only above a given schizophrenia PGRS value, individuals would transition to psychosis and display cognitive alterations. Also, considering genetic evidence on schizophrenia as a by-product of human evolution^46,47^, one could postulate that the schizophrenia PGRS may confer some evolutionary advantages here manifested as increased cognitive performance.

It is worth noting that despite the important neuromaturational events occurring during adolescence that might affect schizophrenia liability (e.g., synaptic pruning and apoptosis)^48^, the results here do not evidence age-related modulation of the PGRS- and ROH-IQ associations. This ostensible independence of genetic effects on cognitive profiles seems to be stable across the considered age range, even when the age-cognition relationship clearly changes during adolescence (e.g., plateaus at ~17 years in Supplementary Figure S4 and Supplementary Figure S8).

To our knowledge, this is the first systematic analysis of interactions between ROH and schizophrenia PGRS. The results show a nominally significant effect that deserves mention: schizophrenia PGRS would be positively correlated with IQ particularly in individuals with a high ROH burden. However, in view of the small size of the effects observed here, validation using larger independent samples is needed.

Limitations of the study include the medical conditions present in some participants, recruited from a hospital; however, as shown above, exclusion of individuals with moderate and severe medical status did not invalidate the findings. Moreover, the focus on a specific age range (8-22 years) and ancestry group has increased specificity, but the findings might not be generalizable to populations from different demographic and genetic backgrounds. Besides, the PGRS findings are nominally significant because of the model selection, and the parameters used to call ROH are relatively robust to confounding due to CNVs^49,50^, but one cannot completely rule out the possibility of artifacts in some of the measurements. Similarly, the procedure adopted here to tune PGRS *p*-value thresholds to find a best-fit model has been previously applied^31-33^, but it renders significance of the main PGRS results only nominally significant in this case. Finally, the relatively small number of participants with MRI scans (*n*=516) could have limited the power of brain-genetics tests considerably.

## ACKNOWLEDGEMENTS

Supported by the Research Council of Norway (204966, 249795, 223273, 213837); the South-Eastern Norway Regional Health Authority (2012-047, 2013-123, 2015-073, 2016-083); European Community’s 7th Framework Programme (602450, IMAGEMEND), and Kristian Gerhard Jebsen Foundation.

## AUTHOR CONTRIBUTIONS

A.C.-P. and L.T.W. conceived and designed the experiments, analyzed the data and wrote the manuscript; A.C.-P., T.K., D.A., N.T.D., T.M., and L.T.W. performed the experiments, and all authors provided substantial input to and gave approval of the final version of the manuscript.

## CONFLICT OF INTEREST

The authors declare no conflict of interest.

